# Differing patterns in community composition and diversity of biofilm-forming bacteria and fungi along a large salinity gradient

**DOI:** 10.64898/2026.10.28.762137

**Authors:** Leena Virta, Jianjun Wang, Iines Salonen

## Abstract

**Aim:** Biofilms are complex ecosystems formed by phototrophic and heterotrophic organisms, such as bacteria and fungi. Biofilms are hotspots for microbial activity and crucial for the functioning of all aquatic ecosystems, but they are overlooked and poorly studied, especially along large environmental gradients. We aimed at resolving the responses of bacterial and fungal communities to environmental variation along a gradient with strong variation in environmental factors, especially salinity. We also investigated whether the diversity of biofilm-forming bacteria and fungi follows the pattern predicted by the ‘Remane curve’, which posits that species diversity is lowest at intermediate salinity.

**Location:** 2300-km-long spatial gradient in the Baltic Sea, Northern Europe Time period: August 2018

**Major taxa studied:** Biofilm-forming bacteria and fungi

**Methods:** We investigated the composition and diversity of bacteria and fungi by collecting biofilm samples from 42 shallow benthic ecosystems. We also characterized key environmental variables, such as salinity, nutrient concentrations, wind exposure, and climatic conditions. Bacterial and fungal communities in biofilm samples were identified using Illumina MiSeq sequencing.

**Results:** We found that bacteria and fungi responded differently to environmental variation. Bacterial communities were mainly controlled by salinity, and their diversity was lowest in the intermediate salinity. Bacterial communities were also dominated by different bacterial orders in high, intermediate, and low salinity. Fungal communities, instead, were jointly controlled by several environmental variables, i.e., salinity, water temperature, wind exposure, and nutrients, and their diversity or composition did not show a clear pattern along the salinity gradient.

**Main conclusions:** To our knowledge, this is one of the first studies to investigate the biofilm-forming communities, especially fungal communities, along large environmental gradients. Our results highlight the role of biofilms as complex ecosystems where organisms’ ecological requirements are unique and variable.

## Introduction

Biofilms, i.e., microbial communities growing attached to biological and non-biological surfaces, are dynamic and structurally complex ecosystems, and one of the oldest, most widely distributed, and most successful forms of life on Earth (Flemming et al. 2016; Westall et al. 2001). Marine biofilms typically consist of different microbial groups, such as bacteria, fungi, archaea, and microalgae, and they perform crucial ecosystem functions and services. For example, they drive biogeochemical cycling processes of several elements, form the basis of food webs, form habitats for other organisms by stabilizing sediments, and improve water quality by removing nutrients (Ortiz-Estrada et al. 2019; Hope et al. 2020). Biofilms are also incredibly productive; 4-11% of the total biomass on Earth has been estimated to constitute of bacteria and archaea in biofilms (Sentenac et al. 2022). For several microbial groups, such as bacteria, biofilms are the main life form (Flemming and Wuertz 2019), and the abundance of microbes in the marine biofilm is several tenfolds higher than in the water column (Qian et al. 2022). However, our knowledge on biofilms in the marine ecosystem is still very limited.

Research on marine biofilms has mostly focused on their negative effects on artificial substrates, such as biofouling and biocorrosion of ships, whereas marine biofilms that grow on natural surfaces and sustain ecosystem functioning have been largely overlooked (Qian et al. 2022). Here, our aim is to decrease this knowledge gap by focusing on biofilms living on natural cobbles in shallow coastal ecosystems in the Baltic Sea.

Microbes in marine biofilms are sensitive to the variation of numerous environmental factors, both local factors, such as nutrients, water temperature, acidity, or turbidity (Qiao et al. 2025), and large-scale variables, such as climate or geographical location (Daufresne et al. 2009; Jyrkänkallio-Mikkola et al. 2017). However, in areas with pronounced salinity gradients, such as estuaries and coastal seas, salinity has been found to override the effect of all other environmental factors in determining microbial diversity and community composition (Herlemann et al. 2011; Klier et al. 2018; Virta et al. 2020). Along a salinity gradient from oligohaline (freshwater) to mesohaline (brackish) and finally polyhaline (marine) waters, species specialized in freshwaters typically dominate the microbial communities in areas with salinity <5 and species specialized in marine waters in areas with salinity >18, whereas communities in brackish areas with salinity 5-18 consist of just a small percentage of species specialized in brackish waters and are rather dominated by freshwater and marine species with large salinity tolerances (Klier et al. 2018; Virta et al. 2020). The low number of species specialized in brackish water may stem from the small total area and isolated nature of brackish water areas in the world and, in the case of the Baltic Sea, the young geological age of the water basin (Olli et al. 2019). For many aquatic organisms, such as macroinvertebrates and phytoplankton, the low number of species specialized in brackish waters and the considerable physiological adaptations that freshwater and marine species face when trying to adapt to brackish waters have led to a pattern of minimal species richness in brackish waters, namely waters with salinity 5-8 (Remane 1934; Olli et al. 2019). However, whether this pattern, called the Remane curve (Artenminimum), also applies to bacteria and fungi living in biofilms is still unexplored.

The Baltic Sea provides an ideal study area for resolving the variation in microbial communities along large environmental gradients that are key for understanding how ecological patterns change in relation to the environment (Thrush et al. 2006). In the Baltic Sea, salinity forms a strong and temporally stable gradient, varying from freshwater (salinity ∼1 in the northernmost part) to almost fully marine (salinity ∼28 in the transition zone from the Baltic Sea to the North Sea). As the salinity gradient stretches over a geographical distance of ∼2300 km, it correlates with latitudinal and climatic gradients, namely from 65.8°N and continental climate in the northern part to 55.7°N and maritime temperate climate in the southern part. Other environmental variables affecting biofilm organisms, such as water nutrient concentration and disturbance by wind and waves, do not form such strong north-south gradients in the Baltic Sea but are highly variable among sites due to the strong but variable anthropogenic effect and varying land-use practices, and the complex nature of the coastline and archipelago. The Baltic Sea may be one of the most studied water basins in the world, but, surprisingly, there is very little knowledge on communities living on very shallow coastal bottoms. These ecosystems are hotspots for benthic productivity and crucial for many ecosystem services as they are situated in the transition zone between land and sea. Biofilms in these illuminated ecosystems are very diverse, because they consist of both phototrophic (e.g., microalgae, cyanobacteria) and heterotrophic (e.g., heterotrophic bacteria, fungi, archaea) organisms. Although the Baltic Sea is tideless and shallow benthic habitats thus always remain submerged, communities living in these environments face strong temporal and spatial environmental variation in terms of, for example, temperature fluctuations, light availability, and ice scouring during winters. Therefore, these communities also face the consequences of climatic and environmental change upfront.

To improve knowledge on organisms in the ecologically crucial but vulnerable shallow coastal ecosystems, we studied biofilm-forming bacteria and fungi living on natural cobbles at 42 shallow sites along the entire environmental gradient of the Baltic Sea. The specific aims were to characterize the community composition and diversity of these communities using Illumina MiSeq sequencing and to resolve the environmental, spatial, and temporal effects on the variation in community metrics. To the best of our knowledge, this is one of the first studies to investigate biofilm-forming organisms on long environmental gradients. We predicted that (i) the effects of environmental and spatial variables would lead to significant changes in the community composition and diversity of bacteria and fungi among environmentally different sampling regions (Lobo et al. 2024), (ii) salinity would override the effects of other environmental variables, and that the diversity of organisms would be lowest in the salinity of 5-8 due to the challenges that brackish waters pose to aquatic organisms (Remane 1934; Virta et al. 2020), (iii) the community composition of the biofilm organisms would be temporally stable during our sampling campaign of three weeks, because environmental conditions remain considerably stable during peak productive season (Virta et al. 2020).

## Materials and methods

### Study area and sampling

Our study area extended over ca. 2300 km along the entire coastline of Sweden from the Bay of Bothnia (northern Baltic Sea; 65.77°N, 22.68°E) to southern Baltic Sea and finally to the transition zone between the Baltic Sea and the North Sea (58.77°N, 11.19°E; Fig. 1). We chose this sampling area because environmental conditions, especially salinity and climate-related factors, form a strong and temporally stable north-south gradient. Salinity increases from ∼28 in the south-west to ∼1 in the north, because the exchange of saline water from the North Sea to the Baltic Sea is restricted and only occurs through the narrow Danish Straits, and because several freshwater rivers discharge into the northern parts of the Baltic Sea. The climate in the south is characterized by mild winters, ice-free seas year-round, and an annual average air temperature of ∼8°C, whereas the climate in the north is defined by cold winters, ice-covered seas during winters, and an annual average air temperature of ∼1°C. We conducted the sampling during peak productive season, on 9th – 24th August 2018. We collected samples from 47 sites and then re-visited the first six sites to study temporal variation. For the selection of study sites, we focused on areas with pronounced changes in environmental conditions, especially salinity. To minimize the effects of land use and other terrestrial factors, we selected sites at far ends of peninsulas or islands, and far away from river mouths. Sampling was conducted following the modified biofilm sampling recommendations by Kelly et al. (1998). At each site, we randomly picked twenty cobble-sized stones along the shoreline from depths of 20-50 cm. We collected the biofilm by scraping the surface of each stone with a sponge (25 cm² per stone) and pooled the accumulated suspension into a composite sample, which was then stored in cold (−18°C) and dark conditions in a portable freezer until further processing. To account for the stability of the substrate, which may affect the growing conditions of biofilm organisms, we measured the volume (length width height) of each sampling stone. We also measured salinity, pH, and water temperature with a portable probe *in-situ* and collected a water sample from each site. These water samples were frozen immediately after sampling and later used for nutrient analyses (NO_2_^−^ + NO_3_^−^, NH_4_^+^, PO_4_^3−^, and Si).

**Fig. 1.**
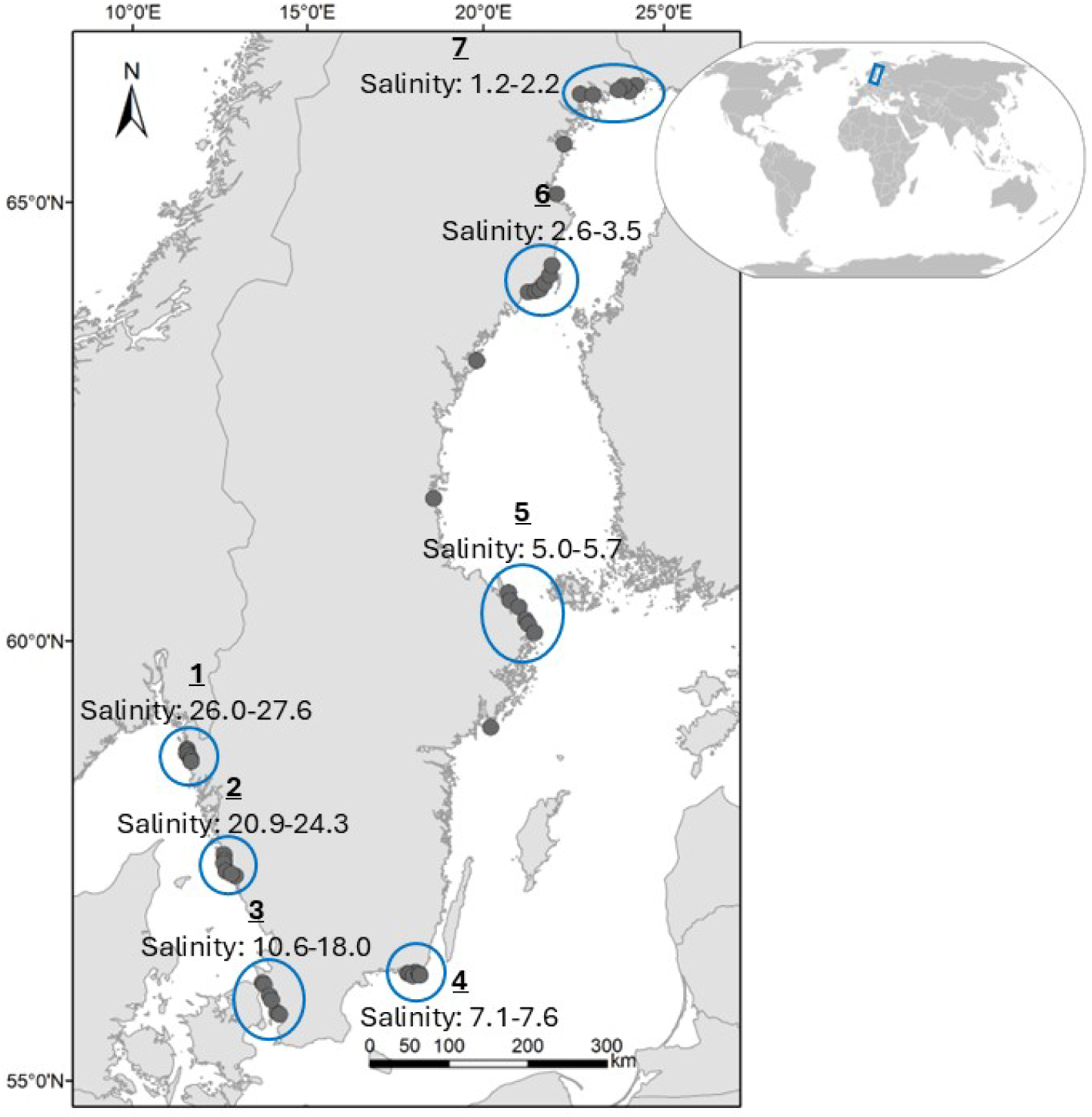
Map of the sampling gradient along the coast of Sweden in the Baltic Sea, Northern Europe. Sampling focused on seven circled areas with different salinity levels.

### Nutrient analyses, wind exposure, and climatic variables

Water nutrient analyses were conducted with an automated photometric analyzer (Thermo Scientific Aquakem 250 [Thermo Fisher Scientific Oy, Vantaa, Finland]) for NO_2_^−^ + NO_3_^−^, PO_4_^3−^, and Si, or manually for NH_4_^+^. To account for the wind and wave exposure of the sampling sites, we calculated fetch, i.e., the distance over which wind can travel across open water, using the ruler tool in Google Earth (Google). We placed the beginning of the ruler exactly at each study site and measured the distance to the next continental shore, island, or islet along 36 lines, which were 10° apart from each other. Thus, we obtained 36 values for each site and used the sum of these values as the measure of the fetch of the sampling site (Mason et al. 2018). To account for climate, we extracted the average annual temperature and average annual precipitation (representative of 1950-2000, spatial resolution ∼ 1km²) from WorldClim database (Hijmans et al. 2005) and calculated climatic values for each sampling site with the ArcGIS 10.2.1, Arcmap-application.

### Genomic DNA extraction and amplicon sequencing

Genomic DNA was extracted from biofilm samples using the PowerSoil DNA Isolation Kit (MoBio, Carlsbad, USA). The bacterial 16S rRNA gene was amplified using modified primers 515F (5′-GTG YCA GCM GCC GCG GTA A-3′, Parada et al., 2016) and 806R (5′-GGA CTA CNV GGG TWT CTA AT-3′, Apprill et al. 2015), targeting the V4 region. Fungal ITS2 regions were amplified with primers gITS7F (5′-GTG ART CAT CGA RTC TTT G-3′) and ITS4R (5′-TCC TCC GCT TAT TGA TAT GC-3′) (Ihrmark et al. 2012). PCR amplification was conducted in 50 μL reaction volumes containing 1 μL of Phanta Max Super-Fidelity DNA Polymerase (Vazyme, China), 25 μL of 2× Phanta Max Buffer, 1 μL of dNTP Mix (10 μM each), 2 μL each of forward and reverse primers, 1 μL of template DNA (20–30 ng/μL), and nuclease-free water to volume. The thermal cycling program for 16S rRNA gene amplification consisted of an initial denaturation at 95°C for 3 min; followed by 30 cycles of 95°C for 10 s, 60°C for 10 s, and 72°C for 45 s; with a final extension at 72°C for 5 min. Fungal ITS2 amplification followed the same conditions, except that the annealing temperature was set at 52°C and 35 cycles were performed. PCR products (2 × 250 bp) from triplicate reactions were pooled, purified, and sequenced on the Illumina NovaSeq 6000 platform (Illumina, USA). The detailed methods can be found in Zhao et al. (2024).

### Bioinformatical and statistical analysis

Sequence data was trimmed, filtered, merged and converted into amplicon sequence variant (ASV) tables using the Chipster software (Kallio et al. 2011) following the in-built Dada2-pipeline (Callahan et al., 2016). Taxonomy was assigned either against the Silva 138.1 database (bacterial sequences, Quast et al. 2023) or the UNITE v. 10 database (fungal ITS sequences, Abarenkov et al. 2024). ASV-tables, taxonomic assignment and environmental metadata were combined into a phyloseq-object (version 1.50.0, McMurdie and Holmes 2013) for further analysis in R version 4.4.3 (R core team 2025). To correct for differences in the sampling depth (library size), cumulative sum scaling (CSS) was applied for median-like quantile normalization (Paulson et al. 2013).

We studied statistical dependence between environmental explanatory variables with Spearman’s rank correlation coefficients and considered values r_s_ > |0.7| as strong correlations. To analyze the statistical significance of the environmental factors, we applied permutational multivariate analysis of variance (PERMANOVA) on the Bray-Curtis dissimilarity matrix of ASV abundances of bacterial or fungal communities as the dependent variable using the adonis2 function in the R package vegan (version 2.6.8, Oksanen et al. 2024). Shannon diversity was calculated using the estimate richness function in phyloseq (McMurdie and Holmes 2013). Pearson correlations between the significant environmental factors and the 10 most abundant bacterial and fungal orders were analyzed using the R package microViz (version 0.12.6, Barnett et al. 2021).

## Results

Environmental variables were highly variable along the sampling gradient. Salinity (1.2-27.6), annual air temperature (1.2-8.1°C), and annual precipitation (482-793 mm) formed increasing gradients from north to south. The variation of other environmental variables was more random among sites. Water temperature varied between 10°C and 25.3°C, pH between 6.7 and 9.5, NH_4_^+^ between 1.4 µg/l and 74.2 µg/l, NO_2_^−^ + NO_3_^−^ between 1.6 µg/l and 64.4 µg/l, PO_4_^3−^ between 1.7 µg/l and 632.8 µg/l, Si between 43.9 µg/l and 1757.3 µg/l, and wind exposure (i.e., fetch) between 140 m and 43725700 m. According to Spearman’s rank correlation, Si, air temperature, precipitation, latitude, and longitude were highly correlated (r_s_ > |0.7|) with salinity. Because we expected salinity to be the overriding factor for the distribution of benthic organisms along our sampling gradient, we chose to keep salinity in further analyses and exclude the other correlating variables.

### The effects of environmental factors on bacterial and fungal community composition and diversity

The community composition of bacteria was significantly controlled by salinity and water temperature (Table 1; Fig. 2). Along the salinity gradient, the bacterial diversity was highest in high (> 20.9) and low (< 2.2) salinity and lowest in intermediate (5-5.7) salinity (Fig. 3).

**Fig. 2.**
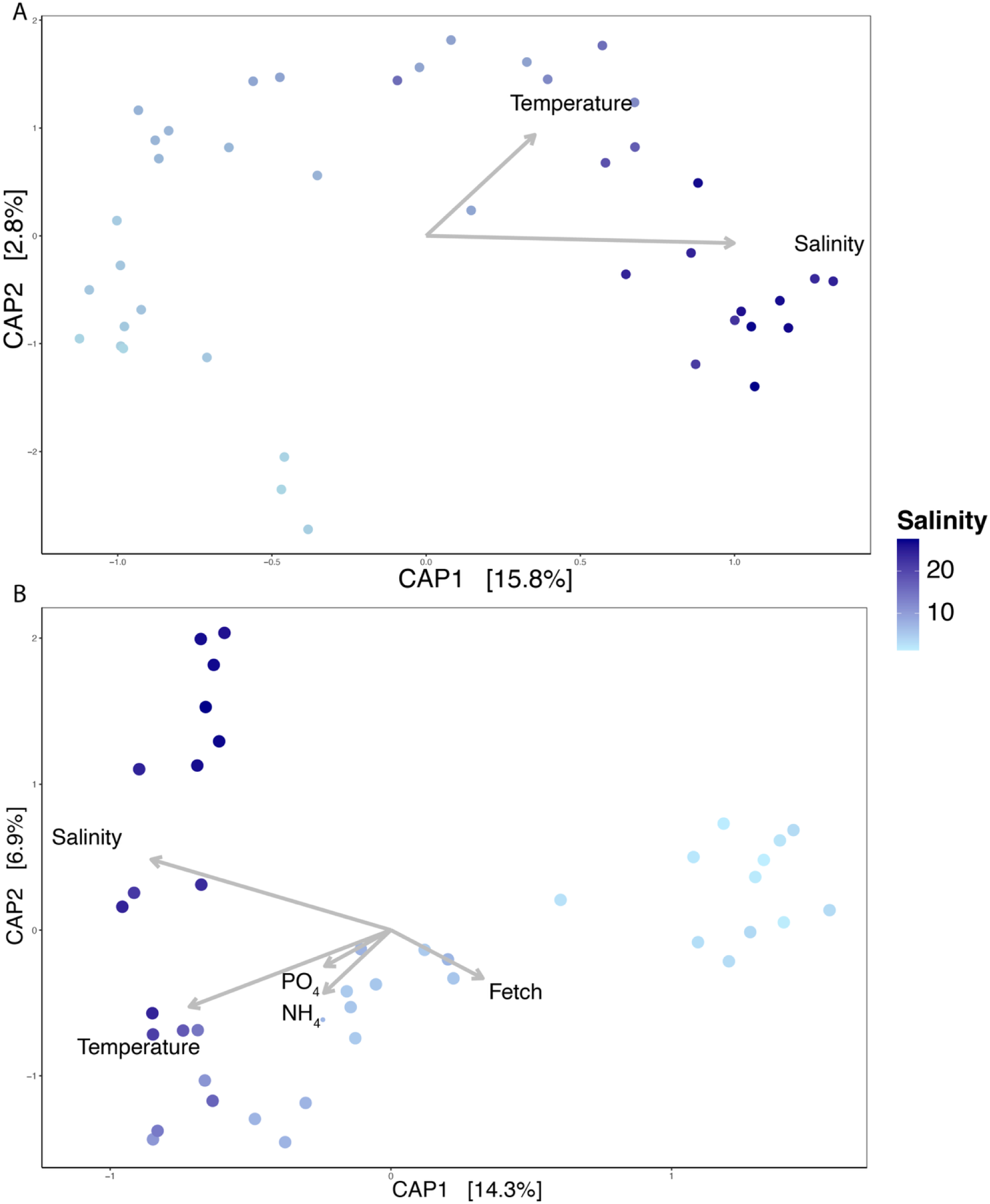
Canonical analysis of principal coordinates (CAP) to illustrate the association of environmental variables on changes in the A) bacterial and B) fungal community composition. Grey arrows represent significant environmental factors and dots each sample along the salinity gradient.

**Fig. 3.**
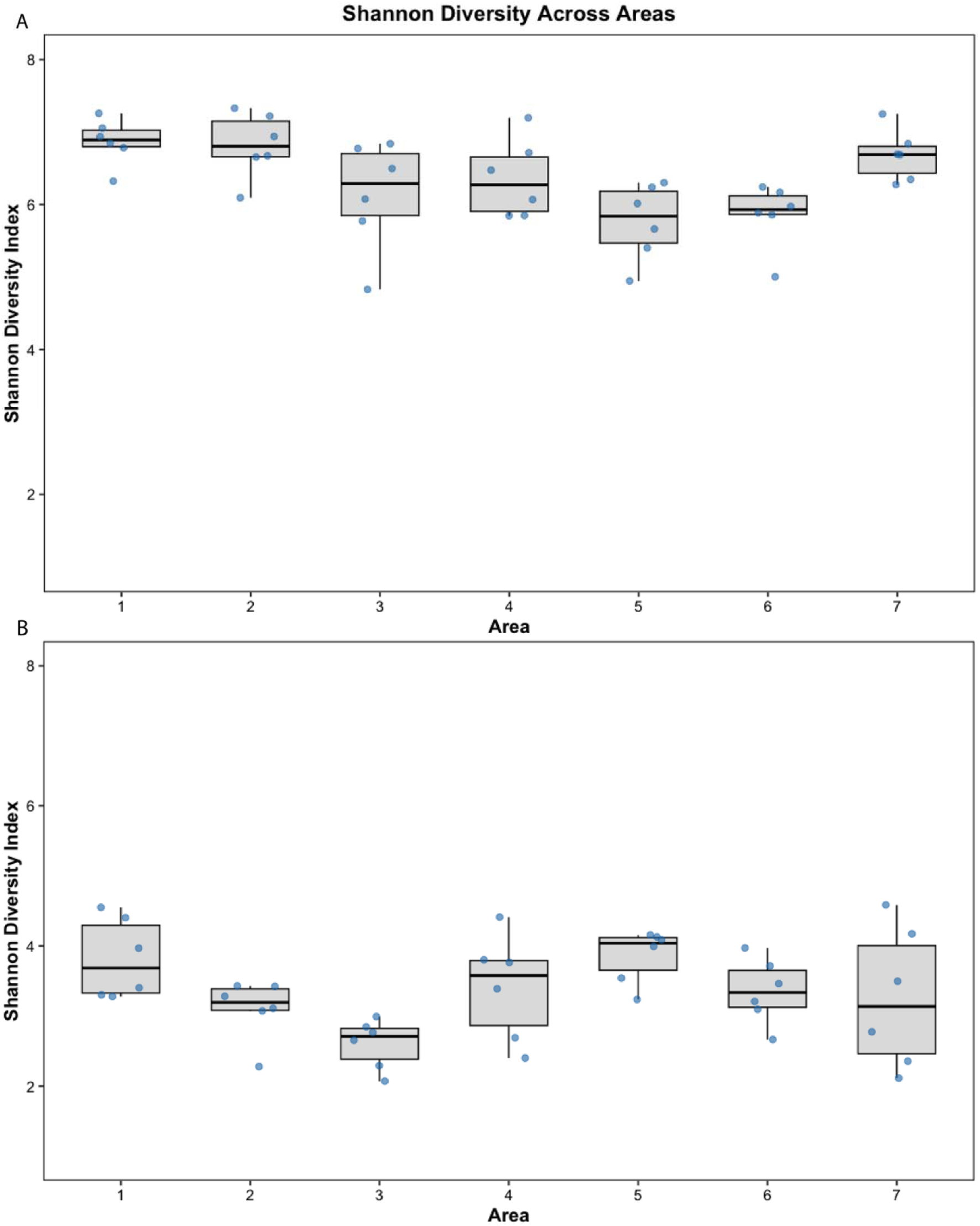
Box plots showing the Shannon’s diversity of a) bacterial and b) fungal communities along the salinity gradient.

**Table 1.** Results of the PERMANOVA analyses for the effects of environmental variables and time (sampling area 5) on the community compositions of bacteria and fungi. Fetch denotes wind exposure. < 0.001 ***, < 0.01 **, < 0.05 *

|  | **Bacteria** |  | **Fungi** |  |
| --- | --- | --- | --- | --- |
|  | <u>p-value</u> | <u>Significance</u> | <u>p-value</u> | <u>Significance</u> |
| Salinity | 0.001 | *** | 0.001 | *** |
| Temperature | 0.013 | * | 0.001 | *** |
| Fetch | 0.148 |  | 0.005 | ** |
| NH<sub>4</sub><sup>+</sup> | 0.754 |  | 0.016 | * |
| PO<sub>4</sub><sup>3-</sup> | 0.182 |  | 0.027 | * |
| NO<sub>2</sub><sup>-</sup> + NO<sub>3</sub><sup>-</sup> | 0.159 |  | 0.690 |  |
| pH | 0.575 |  | 0.861 |  |
| Time | 0.003 | ** | 0.444 |  |

The community composition of fungi was significantly controlled by salinity, water temperature, wind exposure, and the availability of PO_4_^3−^ and NH_4_^+^ (Table 1; Fig. 2). The fungal diversity was variable and showed no trend along the salinity gradient (Fig. 3). The diversity was highest in intermediate (5-5.7) and high (> 26) salinity and lowest in higher intermediate (10.6-18) and low (< 2.2) salinity. There was also considerable variation in fungal diversity within each individual salinity level.

### Variation in the composition of bacterial and fungal communities along environmental gradient

The composition of bacterial communities varied across different salinity levels (Fig. 4). At the order level, the abundance of *Chitinophagales*, *Enterobacterales*, *Flavobacterales*, and *Phormidesmiales* decreased from high salinity to low salinity, whereas the abundance of *Exiguobacterales*, *Cyanobacterales*, and *Cytophagales* increased towards low salinity. Brackish sampling sites (salinity ∼5-7.5) were characterized by high abundance of *Exiguobacterales* and *Micrococcales*. Some orders that were found in low abundance were characteristic to sites with specific salinity levels, such as *Caulobacterales* (salinity 18.9-24.8), *Campylobacterales* (salinity 19.1-22.7), *Rhodothermales* (salinity 22.2-24.8), *Thiotrichales* (salinity 10.6-18), and *Leptolyngbyales* (salinity 1.2-2.2). Pearson’s correlation showed that there was variation in the response of the ten most abundant bacterial orders to salinity and temperature gradients (Fig. 5). The abundance of *Cytophagales*, *Cyanobacterales*, and *Sphingomonadales* correlated negatively with salinity and temperature, whereas the abundance of *Rhodobacterales*, *Flavobacterales*, *Enterobacterales*, and *Phormidesmiales* correlated positively with salinity and temperature. The abundance of *Pseudomonadales* correlated negatively with salinity and positively with temperature, whereas *Chitinophagales* vice versa. *Exiguobacterales* correlated negatively with salinity. In our NMDS analysis on bacterial communities, community composition formed a gradient along the salinity gradient on axis 1, except three low salinity sites that joined sites with intermediate salinity in the middle of the axis (Fig. 7). On axis 2, communities were further divided into groups according to their sampling areas, reflecting the effect of other environmental variables than salinity.

**Fig. 4.**
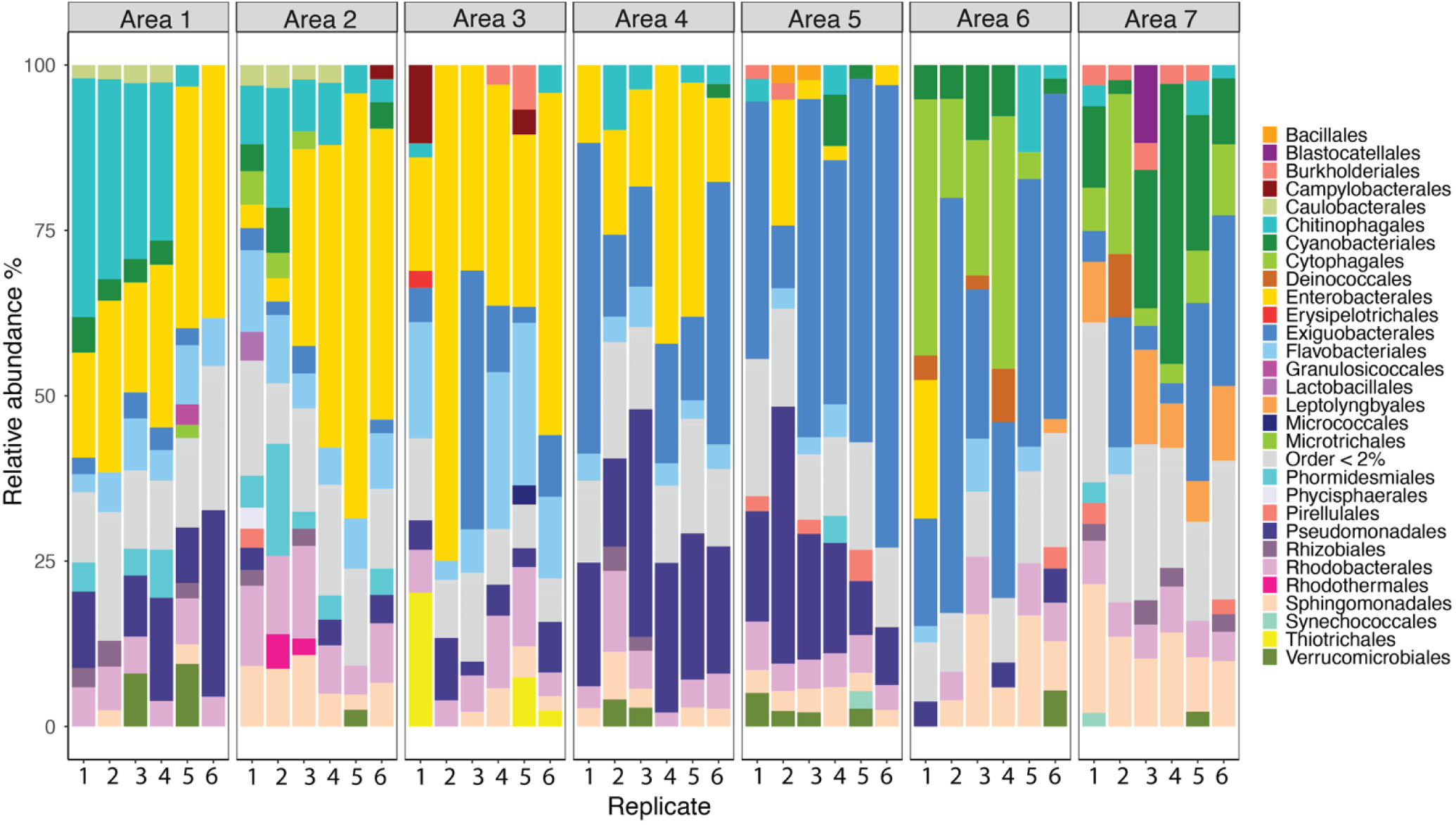
Relative abundance of bacterial orders (RA > 2%) in biofilms of sampling areas 1-7. Bacterial orders that are present < 2% RA are grouped in “Order < 2%”.

**Fig. 5.**
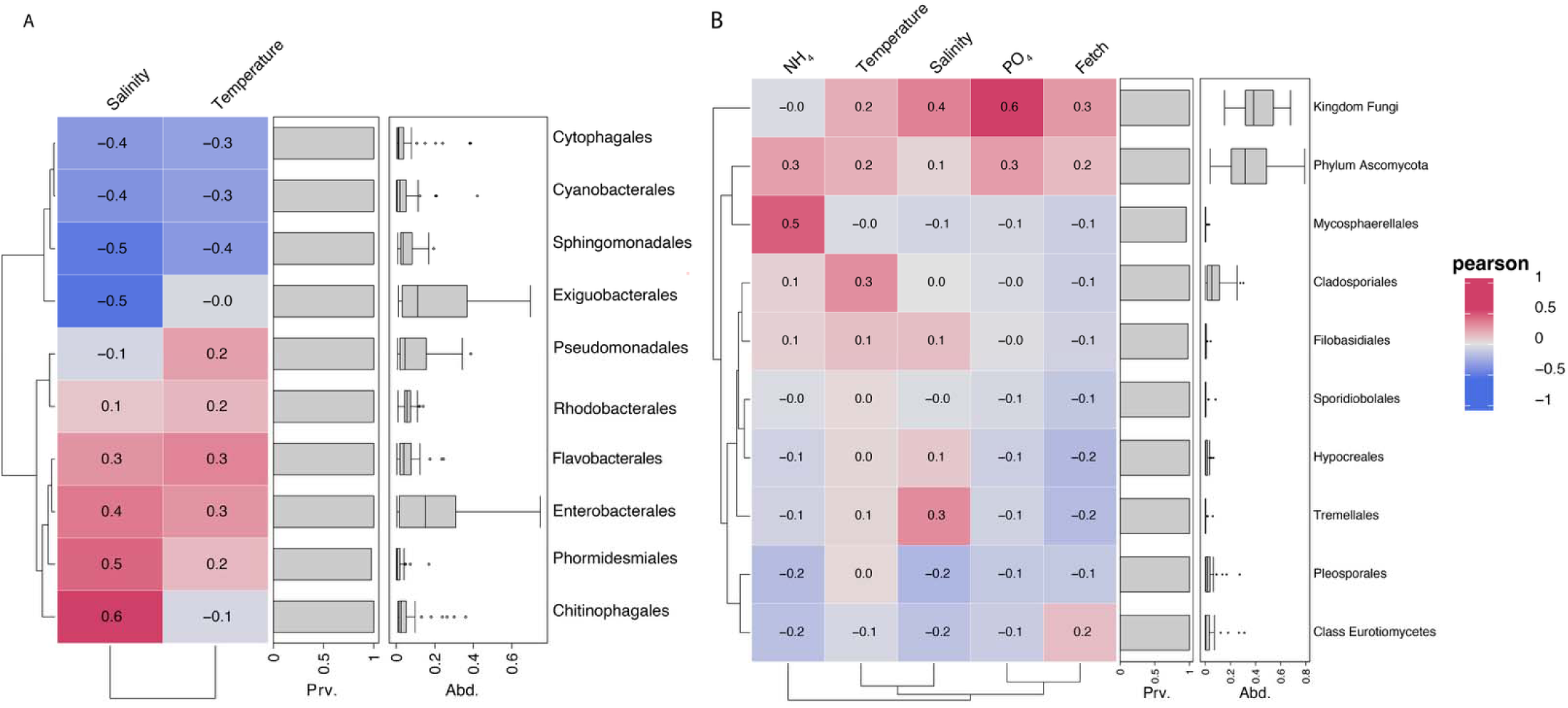
Pearson’s correlations between the most abundant A) bacterial and B) fungal orders and significant environmental variables indicated by PERMANOVA.

The community composition of fungi contained more unidentified taxonomy than bacteria, as ∼50% of sequences at each site could only be identified to domain level (Fig. 6). Shannon diversity (H’) of fungi ranged from 2.17 to 4.58, with the average of 3.34 (±0.69) and was therefore considerably lower than that of bacteria, which it ranged from 4.83 to 7.33 with the average of 6.35 (±0.63). At the phylum level, fungal communities in all salinities were dominated by *Ascomycota*. At the order level, *Cladosporiales* were abundant in the intermediate salinity, present but less abundant in the high salinity, and absent in the low salinity. Pearson’s correlations showed that the seven most abundant fungal taxa that we were able to identify to order level (*Mycosphaerellales*, *Cladosporiales*, *Filobasidiales*, *Sporidiobolales*, *Hypocreales*, *Tremellales*, and *Pleosporales*) responded differently to environmental variables (Fig. 5). The abundances of *Filobasidiales*, *Hypocreales*, and *Tremellales* correlated positively and the abundances of *Mycosphaerellales* and *Pleosporales* negatively with salinity. The abundances of *Cladosporiales*, *Filobasidiales*, and *Tremellales* correlated positively with temperature. The abundances of *Mycosphaerellales* correlated positively and abundances of *Pleosporales* negatively with ammonium abundances. The abundances of all seven orders correlated negatively with wind exposure and PO_4_^3−^. The composition of fungal communities changed gradually along the salinity gradient, as shown on the first axis of NMDS (Fig. 7).

**Fig. 6.**
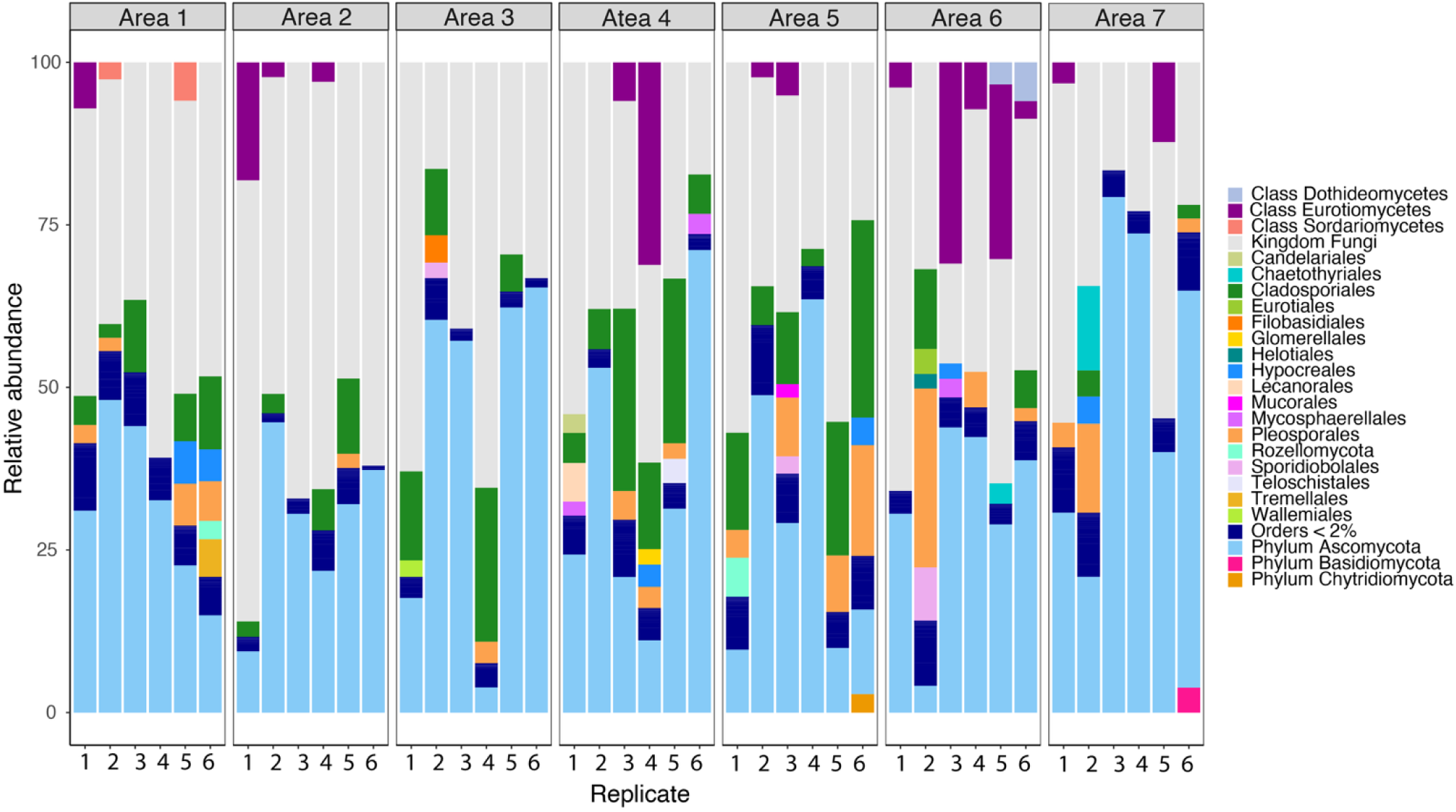
Relative abundance of fungal orders (RA > 2%) in biofilms of sampling areas 1-7. Sequences that could only be identified to class level are marked with c, to phylum level with p, and domain level with k. Fungal orders that are present < 2% RA are grouped in “Order < 2%”.

**Fig. 7.**
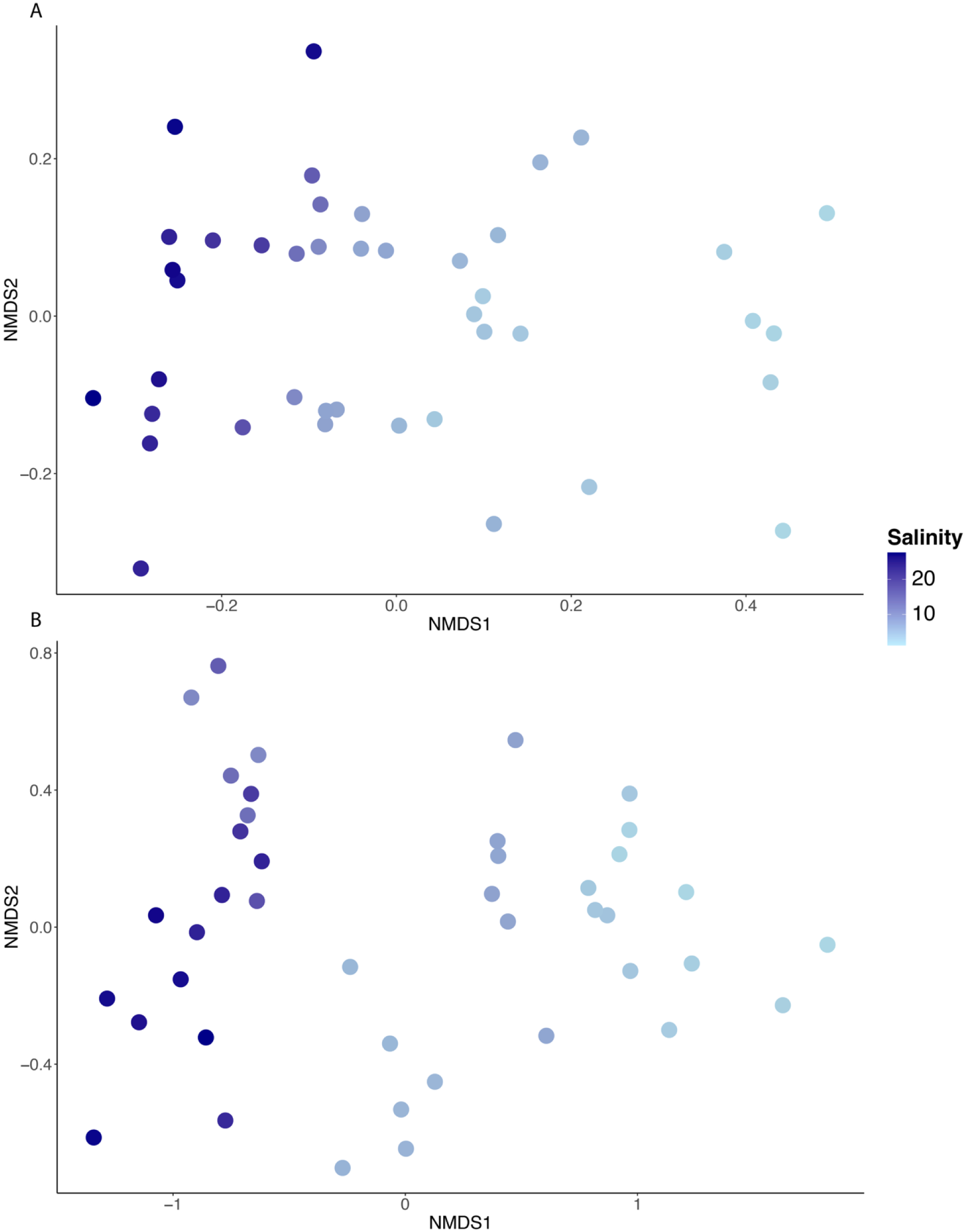
Illustrations for Nonmetric Multidimensional Scaling (NMDS) for a) bacterial and b) fungal community compositions along the salinity gradient.

### Temporal variation of bacterial and fungal community composition

We further investigated the temporal variation of communities on six sites at intermediate salinity from August 9^th^-10^th^ to August 23^rd^-24^th^. Although the composition of bacterial orders remained almost similar, time was a significant factor influencing the relative abundances of the bacterial communities (Table 1; Fig. 8). During this time, the abundance of *Enterobacterales* and *Burkholderiales* increased, whereas the abundance of *Exioguobacterales* and *Pseudomonadales* decreased. Fungal communities did not show clear temporal changes (Table 1; Figure S1).

**Fig. 8.**
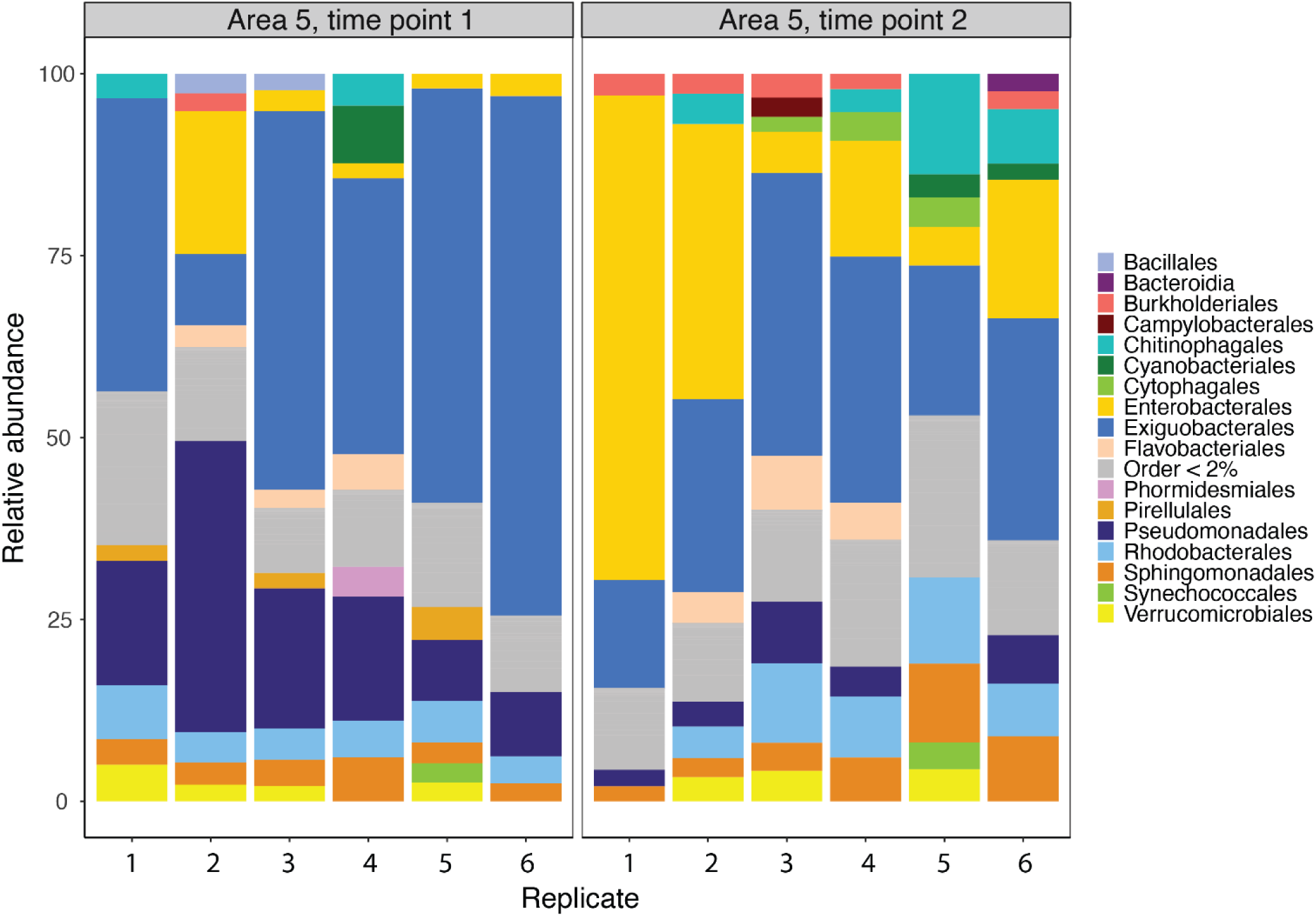
Temporal change in the relative abundance of bacterial orders (RA > 2%) in biofilm samples on the six sites of sampling area 5 during three weeks in August 2019. Orders that are present < 2% RA are grouped in “Order < 2%”.

## Discussion

Biofilms are extremely productive habitats and key to many ecosystem functions. However, they are often overlooked in marine research and, hence, our knowledge on organisms living in biofilms and what environmental factors influence them is still poor. We studied bacterial and fungal communities living as biofilms on shallow benthic habitats along the large environmental and spatial gradient of the Baltic Sea. To the best of our knowledge, this is one of the first studies to investigate coastal, shallow water biofilms with comprehensive sampling along a gradient with considerable environmental variation.

### The effect of environmental factors on bacterial and fungal community composition and diversity

The responses of microbial communities to variation in environmental variables were different between bacteria and fungi. The composition of bacterial communities was mainly controlled by salinity, and as we hypothesized, the bacterial diversity showed a clear trend on the salinity gradient, being highest in marine and freshwater and lowest in brackish water. This is interesting, because it suggests that the pattern presented by Remane (1934), i.e., the lowest species richness of organisms in intermediate salinities of 5-8, also applies to biofilm-forming bacteria. This pattern has previously been confirmed for at least macroinvertebrates (Remane 1934), phytoplankton (Olli et al. 2019), zooplankton (Luceno-Moya and Duggan 2017), and biofilm-forming diatoms (Virta et al. 2020), whereas the diversity of bacteria and protists living as plankton have been shown to be indifferent to salinity variation along a gradient from freshwater to brackish and marine water (Herlemann et al. 2011), or even to have a species richness maximum at intermediate salinities (Filker et al. 2018). Thus, it seems that the response of the diversity of biofilm-forming bacteria to salinity resemble more the responses of eukaryotic organisms than those of pelagic bacteria. We suggest that this may be due to the nature of biofilms as compact communities where different autonomous organisms, such as bacteria, fungi, microalgae, and archaea, are in constant interaction with each other, forming a multicellular entity. Thus, biological interaction in the form of, e.g., competition, mutualism, and parasitism, can modify the responses of individual organism groups in different ways than in environments where biological interaction is less intense, such as in the pelagic ecosystem. This finding highlights the importance of considering biological interactions when studying biofilms.

The composition of fungal communities was controlled by salinity but not as strongly as the bacterial communities, and was also significantly affected by as water temperature, wind exposure, and nutrient concentrations. The effect of wind and waves on the composition of benthic communities is due to the physical disturbance they induce on sessile organisms, such as biofilms, which often leads to lower diversity and production of communities at sites with strong wind disturbance (Reed et al. 2011). The significant effect of nutrients on the fungal community composition is probably due to the important role of many fungal species in nutrient cycling (Lobo et al. 2023). Contrary to our hypothesis, fungal diversity did not show a clear pattern on the salinity gradient. Thus, the Remane curve does not seem to apply to biofilm-forming fungi, which agrees with previous studies on benthic fungi in deeper sediments (Lobo et al. 2023).

### Variation in the composition of bacterial and fungal communities along environmental gradient

As we hypothesized, the community composition of bacteria was different among environmentally different sampling areas, and communities formed a continuum on the salinity gradient. Communities in high salinity were characterized by high abundances of *Chitinophagales*, *Enterobacterales*, *Flavobacterales*, and *Phormidesmiales*. However, there is no earlier evidence of the salinity tolerance levels of many of these bacterial groups, and we assume that the distribution of different bacterial orders may also have been controlled by some unmeasured environmental variables, such as organic matter content. For example, *Chitinophagales* are known for their ability to degrade complex organic matter, such as chitin, and they are often abundant in environments with high chitin content (De Corato 2021). The group of *Enterobacterales* is notorious for containing several pathogens to humans, other animals, and plants, and being resistant to antibiotics (De Oliveira et al. 2020). However, they are also common in aquatic environments from freshwater to marine environments and can be associated with different kinds of aquatic organisms, such as algae, mussels, and fish (Janda and Abbot 2021; Galarza-Verkovitch et al. 2023). *Flavobacterales* are abundant in the marine environment, both in the epipelagic zone and as biofilms (Muck et al. 2019; Ortiz-Estrada et al. 2019). In biofilms, their success is related to their ability to move by gliding, which allows them to search for optimal levels of, e.g., light intensity, salinity, temperature, and oxygen (Nagaraj et al. 2017).

Intermediate salinities were characterized by high abundance of *Exiguobacterales* and *Micrococcales*. *Exiguobacterales* are a typical biofilm forming bacterial group (Ma et al. 2023), and both *Exiguobacterales* and *Micrococcales* tolerate wide environmental variation, for example all salinities from freshwater to marine waters (Liu et al. 2020; Pantoja-Agreda and Pajares 2024). This suggests that biofilm-forming bacterial communities, just like the communities of larger organisms, in brackish waters are not dominated by orders or species specialized in brackish water but rather orders or species that tolerate wide variation in salinity (Remane 1934).

Bacterial communities in areas with low salinity were dominated by *Exiguobacterales*, *Cyanobacterales*, and *Cytophagales*. The high abundance of the photosynthesizing *Cyanobacterales* in these low salinity areas is interesting, as it may suggest that cyanobacteria strongly contribute to primary production in biofilms in the freshwater areas of the Baltic Sea, in addition to diatoms. This result agrees with some previous studies that have found diatoms to be more halotolerant than cyanobacteria that prefer lower salinity levels (Khatoon et al. 2010). However, we also measured lowest water temperatures in areas with lowest salinity levels, and the contribution of cyanobacteria has previously been shown to decrease towards lower temperatures (Watermann et al. 1999). This suggests that salinity may be more important than temperature as an environmental factor controlling the contributions of cyanobacteria and benthic diatoms as primary producers in biofilms, at least if variation in salinity along the study gradient is large enough.

However, the abundances of several of the most abundant bacterial orders were also either negatively or positively correlated with water temperature. Our findings agree with previous investigations, because orders whose abundances negatively correlated with temperature were orders that are known to strive in cold and extreme environments, such as *Cytophagales* (Bowman et al. 2000) and *Sphingomonadales* (Faragó et al. 2024), whereas orders whose abundances positively correlated with temperature were orders that are known to strive in elevated temperatures, such as *Rhodobacterales* (Sun et al. 2022).

For fungi, a rather low percentage, i.e., ca. 30-80% of the communities could be taxonomically identified to order or even phylum level, which is consistent with other recent studies on fungal communities in the Baltic Sea (Lobo et al. 2024). The low number of identified orders is probably due to deficiencies in reference libraries, and this shows how little we still know about marine fungi and highlights the importance of studies like this for improving knowledge on this important organism group. The community composition of fungi still formed a gradient along the salinity gradient, thus revealing the strong effect of salinity on fungal communities (Lobo et al. 2024). On phylum level, communities at all sites were dominated by *Ascomycota*. On order level, the most prominent difference among salinity levels were differences in the abundance of *Cladosporiales*. They were highly abundant in intermediate salinities, present but sparse in high salinities, and completely absent in the lowest salinities. *Cladosporiales* are a group of fungi that is ubiquitous in the environment and includes some highly adaptable species and, thus, can adapt to extreme conditions (Peng et al. 2021). We have found no previous evidence on the proclivity of *Cladosporiales* for brackish waters and, therefore, we assume that the high abundance of *Cladosporiales* in waters with intermediate salinity is rather a result of their adaptability to challenging conditions.

Pearson’s correlations showed that the abundance of many fungal orders correlated either positively or negatively with salinity, water temperature, and wind exposure, although most of the correlations were weak. The order with the strongest positive correlation with salinity was *Tremellales*. This could be expected, because *Tremellales* has been previously identified from fully marine environments (Zhu et al. 2023). The order with the strongest positive correlation with water temperature was *Cladosporiales*. This order has previously been identified from both tropical and arctic environments (Laichmanová 2023; Papan et al. 2023), suggesting a wide temperature tolerance. The abundances of all fungal orders correlated negatively with wind exposure. This is due to the disturbing effect of wind-induced waves that makes it difficult for sessile organisms in shallow benthic habitats to stay attached to the substrate. The effect of wind and waves has been shown for many different organisms (e.g., macroalgae in Jonsson et al. 2006; benthic diatoms in Virta et al. 2020) and now benthic fungi, thus we were surprised to not see a significant effect of wind exposure on the diversity of bacterial communities.

### Temporal variation of bacterial and fungal community composition

The composition of bacterial communities changed during our sampling campaign of three weeks during peak productive season in August. Observed bacterial orders remained almost the same, but their abundances changed. Such quick changes in community composition indicate that the bacterial communities in biofilms are more dynamic than the communities of diatoms and other eukaryotes (Korhonen et al. 2010; Virta et al. 2020) and seem to change as rapidly over short timescales as do shallow water planktonic bacterial communities (Hatosy et al. 2013; Cram et al. 2015). The strong temporal variation in the communities of biofilm bacteria is most likely due to changes in the environment that can be especially prominent during highly productive seasons and in high latitudes with strong seasonality. Environmental variables that changed most at our temporal study sites during the temporal study of three weeks were water temperature that, on average, decreased from 21.5°C to 18.1°C and different forms of nitrogen that decreased from 13.1 µg/l to 6.3 µg/l (NH_4_^+^) and from 8.7 µg/l to 6.5 µg/l (NO_2_^−^ + NO_3_^−^). Bacterial orders with most notable abundance changes were *Enterobacterales* and *Burkholderiales* whose abundances increased and *Pseudomonadales* and *Exiguobacterales* whose abundances decreased. The increase in the abundance of *Burkholderiales*, that are a commonly observed and often dominant bacterial group in the Baltic Sea and other estuaries (Piwosz et al. 2013), may be due to the decrease in water temperature, as *Burkholderiales* have been found to dominate bacterial communities during cold seasons (Ameryk et al. 2021). The decrease of the abundance of *Pseudomonadales*, a group of putative denitrifiers, may be due to decreasing nutrient concentrations (Liu et al. 2018). However, the increase in the abundance of *Enterobacterales* and the decrease in the abundance of *Exiguobacterales* during decreasing nitrogen concentration was surprising, because *Enterobacterales* are often related to high nutrient concentrations and *Exiguobacterales* to low nutrient concentrations (Liu et al. 2018; Teixeira et al. 2022).

Following our hypothesis, the composition of fungal communities did not show notable temporal changes during the three-week sampling campaign, even when there were considerable temporal changes during this time for water temperature and nitrogen concentration. These environmental variables have been identified as important environmental constraints for fungal communities (Taylor and Cunliffe 2016). This suggests that eukaryotic organisms in biofilms are less dynamic and respond to environmental changes slower than prokaryotic communities.

To conclude, this study improved our knowledge on biofilms that are important ecosystems for the functioning of aquatic systems but poorly studied and often overlooked. We discovered that organisms living in biofilms respond to environmental variation differently than those living in other habitats, such as in the pelagic habitat, and that different organism groups in the biofilm also have different environmental preferences and requirements. Our results highlight the importance of considering biofilms as complex entities, where different organisms perform different functions.

## Supporting information

Supportive information

## Data and code availability statement

DNA sequence data is deposited in NCBIs Sequence Read Archive (SRA) under Bioproject number PRJNA1452985. The 16S rRNA gene sequences have submission number <u>SUB16108633</u> and ITS gene sequences submission number <u>SUB16123723</u>. R code is archived in Zenodo open repository, https://doi.org/10.5281/zenodo.19628549.

