## Supplementary material for "Differing patterns in community composition and diversity of biofilm-forming bacteria and fungi along a large salinity gradient": Supportive information

**Supportive information for:**

**Communities of biofilm-forming bacteria and fungi have differing patterns of community composition and diversity along a large salinity gradient in the Baltic Sea**

Leena Virta^1,2^, Jianjun Wang^3^ & Iines Salonen^2*^

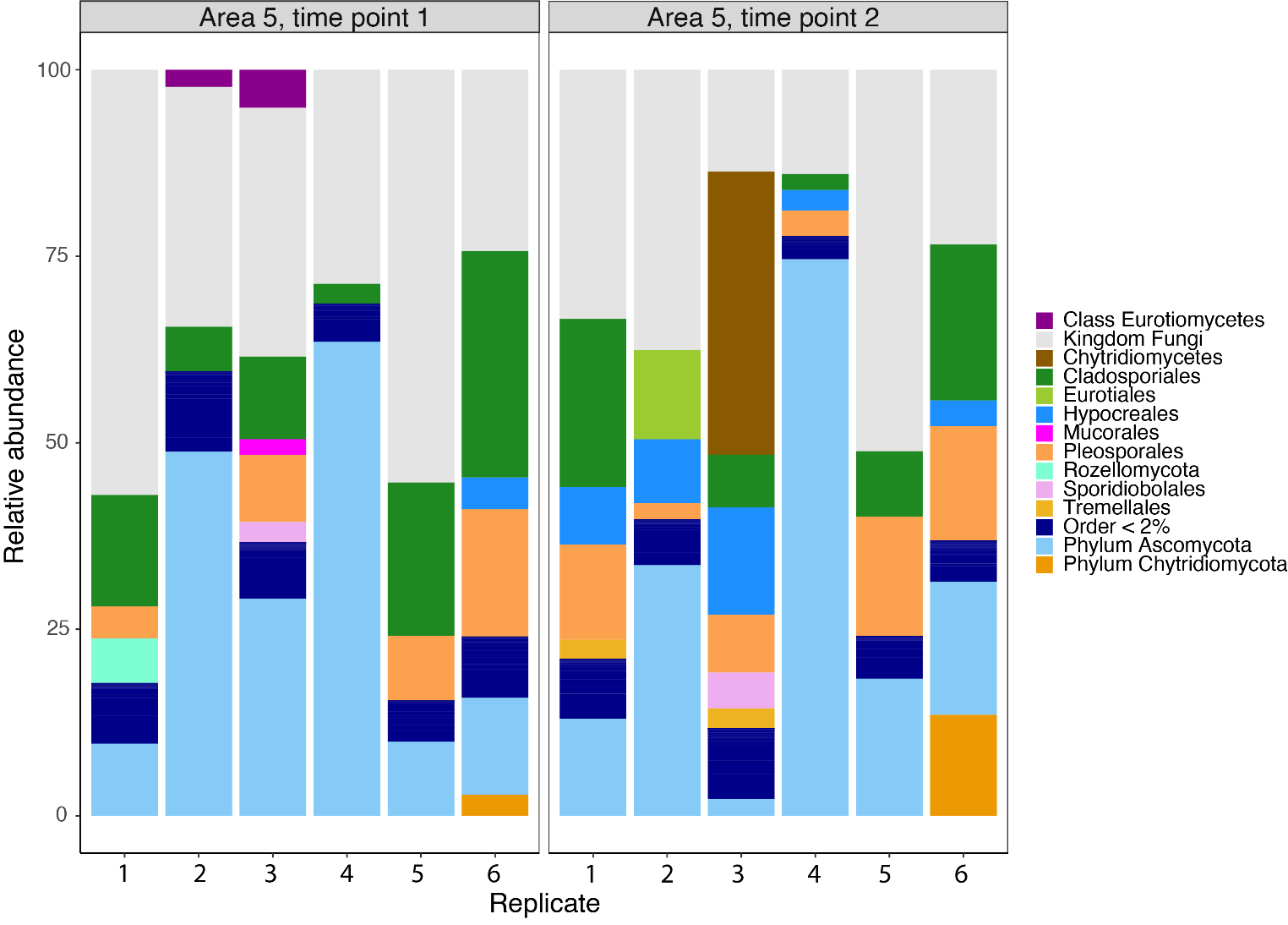

Fig. S1. Temporal change in the relative abundance of fungal orders (RA > 2%) in biofilms on the six sites of sampling area 5 during three weeks in August 2019. Orders that are present < 2% RA are grouped in “Order < 2%”.
